# MNK1 and MNK2 expression in the human dorsal root and trigeminal ganglion

**DOI:** 10.1101/2023.01.04.522773

**Authors:** Stephanie Shiers, James J. Sahn, Theodore J. Price

## Abstract

Mitogen activated protein kinase interacting kinases (MNK) 1 and 2 are serine/threonine protein kinases that play an important role in translation of mRNAs through their phosphorylation of the RNA 5’-cap binding protein, eukaryotic translation initiation factor (eIF) 4E. These kinases are downstream targets for mitogen activated protein kinases (MAPKs), extracellular activity regulated protein kinase (ERK) and p38. MNKs have been implicated in the sensitization of peripheral nociceptors of the dorsal root and trigeminal ganglion (DRG and TG) using transgenic mouse lines and through the use of specific inhibitors of MNK1 and MNK2. While specific knockout of the *Mknk1* gene suggests that it is the key isoform for regulation of nociceptor excitability and nociceptive behaviors in mice, both *MKNK1* and *MKNK2* genes are expressed in the DRG and TG of mice and humans based on RNA sequencing experiments. Single cell sequencing in mice suggests that *Mknk1* and *Mknk2* may be expressed in different populations of nociceptors. We sought to characterize mRNA expression in human DRG and TG for both MNK1 and MNK2. Our results show that both genes are expressed by nearly all neurons in both human ganglia with expression in other cell types as well. Our findings provide evidence that MNK1 and MNK2 are expressed by human nociceptors and suggest that efforts to pharmacologically target MNKs for pain would likely be translatable due its conserved expression in both species.

## Introduction

MNK1 and 2 are serine/threonine protein kinases in the MAPK family (Buxade et al., 2008; Joshi and Platanias, 2014). They are best known for phosphorylation of the 5’ mRNA Cap-binding protein, eIF4E (Waskiewicz et al., 1999). Phosphorylation of eIF4E controls the translation of a subset of mRNAs that are involved in cancer, inflammation, the response to viruses and neuronal plasticity (Herdy et al., 2012; Gkogkas et al., 2014; Moy et al., 2017; Moy et al., 2018b; Megat et al., 2019). In the nervous system, MNK activity has been associated with the excitability of nociceptors in the peripheral nervous system (PNS) (Moy et al., 2017; Megat et al., 2019; Mihail et al., 2019; Barragan-Iglesias et al., 2020; Jeevakumar et al., 2020) and certain forms of synaptic plasticity in the CNS (Aguilar-Valles et al., 2018; Silva Amorim et al., 2018; Shukla et al., 2021; Chalkiadaki et al., 2022). Recently, several additional phosphorylation targets for MNK were described in the brain, including the synaptic protein Ras GTPase activating protein 1 (SYNGAP1), which likely contributes to some features of MNK signaling that are important for social behavioral phenotypes seen in *Mknk* double knockout mice (Chalkiadaki et al., 2022). Knockout and transgenic studies in male and female mice have implicated eIF4E phosphorylation and MNK1 in several forms of inflammatory pain, neuropathic pain, as well as mouse models of migraine headache (Moy et al., 2017; Moy et al., 2018a; Megat et al., 2019; Barragan-Iglesias et al., 2020; Mody et al., 2020; Shiers et al., 2020b; Lackovic et al., 2022). Pharmacological studies show that small molecule MNK inhibitors can prevent and/or reverse established pain phenotypes in the same animal models that have been studied with germline transgenic or knockout approaches (Megat et al., 2019; Shiers et al., 2020b; Lackovic et al., 2022). The existing mouse data suggests that MNK1 may be the most important pharmacological target for pain (Megat et al., 2019; Shiers et al., 2020b; Lackovic et al., 2022), but human studies are needed to firmly establish whether this is likely to be conserved, or not. As a first step toward this goal, we have assessed *MKNK1* and *MKNK2* expression in human DRG and TG at the RNA level.

Mouse and human single cell and spatial sequencing experiments give some preliminary insight into *MKNK1* and *MKNK2* gene expression in DRG and TG. The work of Zeisel and colleagues suggests that both *Mknk1* and *Mknk2* are expressed by most mouse DRG neurons, but *Mknk1* expression is higher in peptidergic nociceptors while *Mknk2* is more highly expressed by non-peptidergic nociceptors (Zeisel et al., 2018). Findings from the work of Renthal and colleagues supports this conclusion except that *Mknk1* gene expression was not detectable in the non-peptidergic subset of nociceptors in mice (Renthal et al., 2020). In the mouse TG at the single cell level, neither *Mknk1* nor *Mknk2* were as robustly detected as in the mouse DRG, but, again, *Mknk1* was mostly expressed in peptidergic nociceptors and *Mknk2* was found in non-peptidergic nociceptors (Yang et al., 2022). In humans there is not a clear distinction between peptidergic and non-peptidergic nociceptors because almost all human nociceptors express the neuropeptide calcitonin gene related peptide (CGRP, *CALCA* gene), the nerve growth factor receptor (NGF) tyrosine receptor kinase A (TrkA, *NTRK1* gene) and transient receptor potential vanilloid type 1 (TRPV1, *TRPV1* gene) (Shiers et al., 2020a; Shiers et al., 2021; Tavares-Ferreira et al., 2022). This suggests that the gradient in *Mknk1* and *Mknk2* expression between nociceptor populations may not be found in humans. Indeed, spatial sequencing results on human DRG show consistent levels of expression for *MKNK1* and *MKNK2* across human nociceptor subtypes (Tavares-Ferreira et al., 2022) but bulk sequencing experiments consistently demonstrate higher levels of expression for the *MKNK2* gene in human DRG samples (Ray et al., 2018; North et al., 2019; Wangzhou et al., 2020; Ray et al., 2022). In human TG, both *MKNK1* and *MKNK2* are detected in single nuclei from neurons, but expression levels are low and sparse across subtypes (Yang et al., 2022).

Given that MNK is a therapeutic target for pain that emerges from activation of nociceptors in both the body (Yousuf et al., 2021) and the head (Lackovic et al., 2022), and that the existing literature does not clearly delineate expression of these two isoforms in the human DRG or TG, we assessed this directly using RNAscope *in situ* hybridization. Our results show that both MNK isoforms are expressed at the RNA level in greater than 95% of human nociceptors. Inhibition of both MNK isoforms should be considered during efforts to develop pain therapeutics targeting MNK modulation.

## Methods

### DRG and TG Tissue preparation

All human tissue procurement procedures were approved by the Institutional Review Boards at the University of Texas at Dallas. Human lumbar DRGs were collected from organ donors through a collaboration with the Southwest Transplant Alliance. Upon removal from the body, the DRGs were frozen in dry ice and stored in a −80°C freezer. DRG donor information is provided in **Table 1**. Fresh frozen human trigeminal ganglia (TG) were received from the Netherlands Brain Bank and stored in a −80°C freezer. TG donor information is provided in **Table 2**. The frozen human DRGs and TGs were gradually embedded in OCT in a cryomold by adding small volumes of OCT over dry ice to avoid thawing. All tissues were cryostat sectioned at 20μm onto SuperFrost Plus charged slides. Sections were only briefly thawed in order to adhere to the slide but were immediately returned to the −20°C cryostat chamber until completion of sectioning. The slides were then immediately utilized for histology.

**Table 1.** Human DRG tissue Information. Donor information is given for all the samples that were used for RNAscope *in situ* hybridization on DRGs.

| Donor # | Tissue | Sex | Age | Race | Surgery/Cause of Death | Collection Site |
| --- | --- | --- | --- | --- | --- | --- |
| 1 | Lumbar DRG | Female | 44 | Black | Anoxia/Cardiac arrest | Southwest Transplant Alliance |
| 2 | Lumbar DRG | Female | 34 | White, non-Hispanic | Anoxia/Opioid overdose | Southwest Transplant Alliance |
| 3 | Lumbar DRG | Male | 29 | Black | Head trauma/Blunt injury | Southwest Transplant Alliance |

**Table 2.** Human TG tissue Information. Donor information is given for all the samples that were used for RNAscope *in situ* hybridization on TGs.

| Donor # | Tissue | Sex | Age | Collection Site |
| --- | --- | --- | --- | --- |
| 2012-070 | TG | Male | 79 | Netherlands Brain Bank |
| 2013-025 | TG | Female | 84 | Netherlands Brain Bank |
| 2020-034 | TG | Female | 96 | Netherlands Brain Bank |

### DRG RNAscope in situ hybridization

RNAscope *in situ* hybridization multiplex version 1 (Cat 320851, *now discontinued*) was performed as instructed by Advanced Cell Diagnostics (ACD) on human DRG sections. The protease incubation period was optimized as recommended by ACD for the specific lot of protease reagent. A table of all probes used is shown in **Table 3**. Following RNAscope, all slides were cover-slipped with Prolong Gold Antifade mounting medium.

**Table 3.** Summary table of Advanced Cell Diagnostics (ACD) RNAscope probes.

| mRNA | Gene name | ACD Probe Cat No. |
| --- | --- | --- |
| *MKNK1* | MAPK interacting serine/threonine kinase 1 transcript variant 1 | 855481-C1 |
| *MKNK2* | MAPK interacting serine/threonine kinase 2 transcript variant 2 | 855491-C1 |
| *SCN10A* | Sodium Voltage-Gated Channel Alpha Subunit 10; Nav1.8 | 406291-C2 and C3 |

### TG RNAscope in situ hybridization

RNAscope *in situ* hybridization multiplex version 2 (Cat 323100) was performed as instructed by ACD. The protease incubation period was optimized as recommended by ACD for the specific lot of protease reagent. A table of all probes used is shown in **Table 3**. Akoya Cy3 (Cat NEL744001KT, Akoya Biosciences) and Cy5 dyes (Cat NEL745001KT, Akoya Biosciences) were used for probe labeling.

### Tissue Quality Check

All tissues were checked for RNA quality by using a positive control probe cocktail (ACD) which contains probes for high, medium and low-expressing mRNAs that are present in all cells (ubiquitin C > Peptidyl-prolyl cis-trans isomerase B > DNA-directed RNA polymerase II subunit RPB1). All tissues showed signal for all 3 positive control probes. A negative control probe against the bacterial DapB gene (ACD) was used to check for non-specific/background label.

### Image Analysis

DRG and TG sections were imaged on an Olympus FV3000 confocal microscope at 20X magnification. Three 20X images were acquired of each human DRG section, and 3 sections were imaged per human donor. The acquisition parameters were set based on guidelines for the FV3000 provided by Olympus. In particular, the gain was kept at the default setting 1, HV ≤ 600, offset = 4, and laser power ≤ 10%. The raw image files were brightened and contrasted in Olympus CellSens software (v1.18), and then analyzed manually one cell at a time for expression of each gene target. Cell diameters were measured using the polyline tool. Total neuron counts for human samples were acquired by counting all of the probe-labeled neurons and all neurons that were clearly outlined by DAPI (satellite cell) signal and contained lipofuscin in the overlay image.

Large globular structures and/or signal that auto fluoresced in all channels (Cy3 and Cy5) was considered to be background lipofuscin and was not analyzed. Aside from adjusting brightness/contrast, we performed no digital image processing to subtract background. We attempted to optimize automated imaging analysis tools for our purposes, but these tools were designed to work with fresh, low background rodent tissues, not human samples taken from older organ donors. As such, we chose to implement a manual approach in our imaging analysis in which we used our own judgement of the negative/positive controls and target images to assess mRNA label. Images were not analyzed in a blinded fashion.

### Data Analysis

Graphs were generated using GraphPad Prism version 8.4.3 (GraphPad Software, Inc. San Diego, CA USA). Population percentage was calculated for each section, and then averaged for each donor. The total number of neurons assessed between all 3 sections for each donor is indicated on the piecharts. Relative frequency distribution histograms with a Gaussian distribution curve were generated using the diameters of all target-positive neurons.

## Results

We first did RNAscope for *MKNK1* mRNA in combination with *SCN10A*, on human DRG to assess its cellular expression in human sensory neurons (**Fig 1**). We found that *MKNK1* mRNA was abundant in virtually all neurons (88.0-97.2%; average 92.4%) in human DRG (**Fig 1A-C**), and in non-neuronal cells as well. These neurons ranged in size from small-to-large diameter (**Fig 1D**). 94.9% of *SCN10A*+ neurons coexpressed *MKNK1* (**Fig 1E**). *SCN10A* is a nociceptor-specific marker in rodent DRG at the protein and mRNA levels (Akopian et al., 1996; Sangameswaran et al., 1996; Dib-Hajj et al., 1999); therefore, these findings are consistent with the notion that *MKNK1* mRNA is present in human nociceptors.

**Figure 1.**
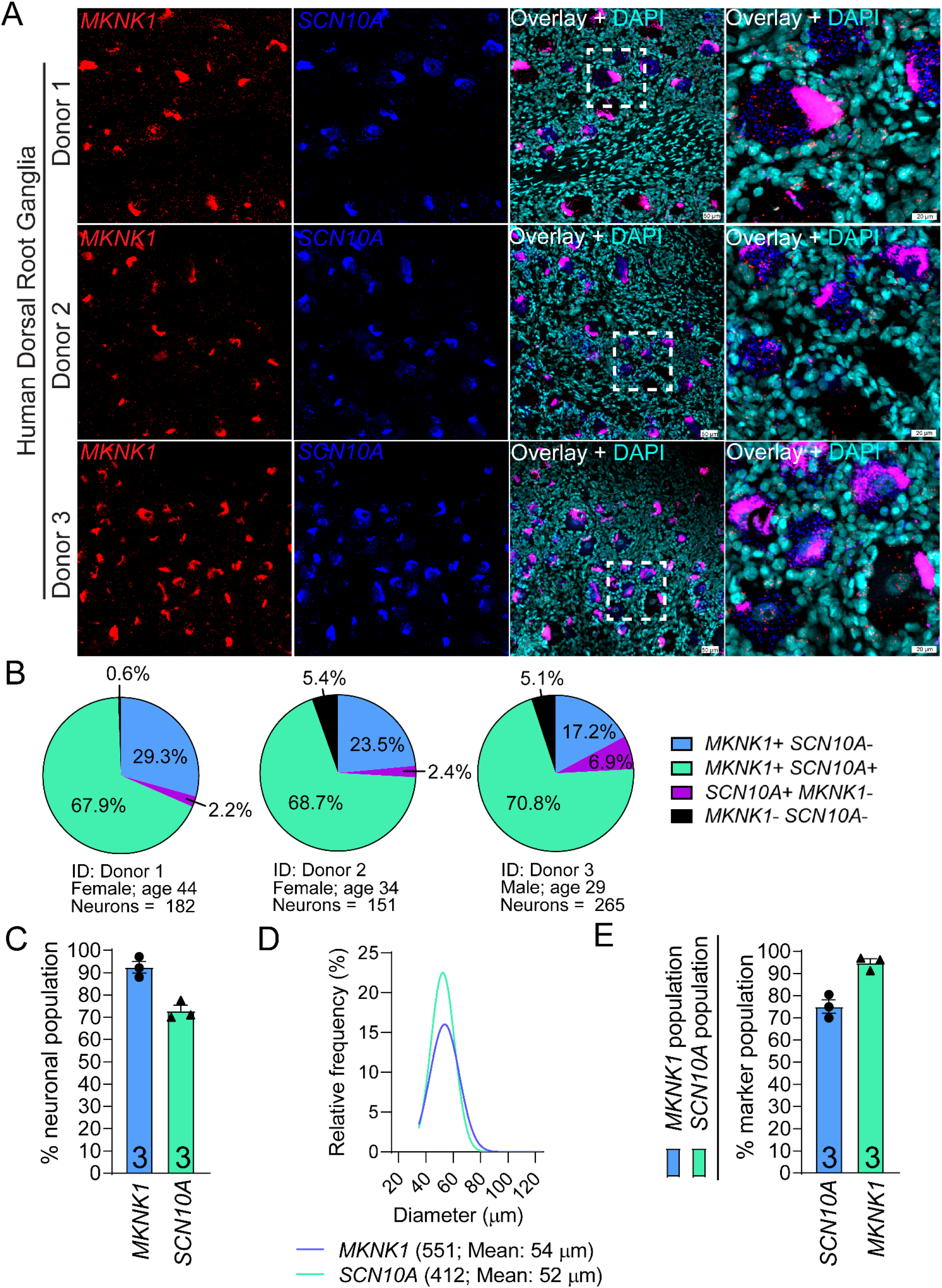
Distribution of *MKNK1* (MNK1) mRNA in human dorsal root ganglia. **A)** Representative 20X images of human lumbar DRGs labeled with RNAscope *in situ* hybridization for *MKNK1* (red) and *SCN10A* (blue) mRNAs and co-stained with DAPI (cyan). The fourth panel for each donor is a zoomed-in region demarcated by white boundaries in the 20X overlay image. Lipofuscin (globular structures) that autofluoresced in both channels and appear magenta in the overlay image were not analyzed as this is background signal that is present in all human nervous tissue. *MKNK1* mRNA was expressed in neurons and non-neuronal cells. **B)** Pie-charts showing the distribution of *MKNK1* neuronal subpopulations in human DRG for each donor. **C)** 92.4% of human DRG sensory neurons were positive for *MKNK1*. **D)** Histogram with Gaussian distribution displaying the size profile of all *MKNK1*+ and *SCN10A*+ neurons in human DRG. **E)** Percentage of *MKNK1*+ neurons that coexpressed *SCN10A* (blue bar), and the percentage of *SCN10A*+ neurons that coexpressed *MKNK1* (green bar). Scale bars: 20X = 50μm. Zoomed in panel = 20μm.

Similarly, we assessed the distribution of *MKNK2* mRNA in human DRG. *MKNK2* mRNA was abundantly expressed in most neurons (90.4-94.0%; average 92.5%) in human DRG (**Fig 2A-C**), and in non-neuronal cells as well. These neurons ranged in size from small-to-large diameter (**Fig 2D**). Like *MKNK1*, most *SCN10A*+ neurons co-expressed *MKNK2* (95.8%) suggesting that *MKNK2* is present in human nociceptors (**Fig 2E**).

**Figure 2.**
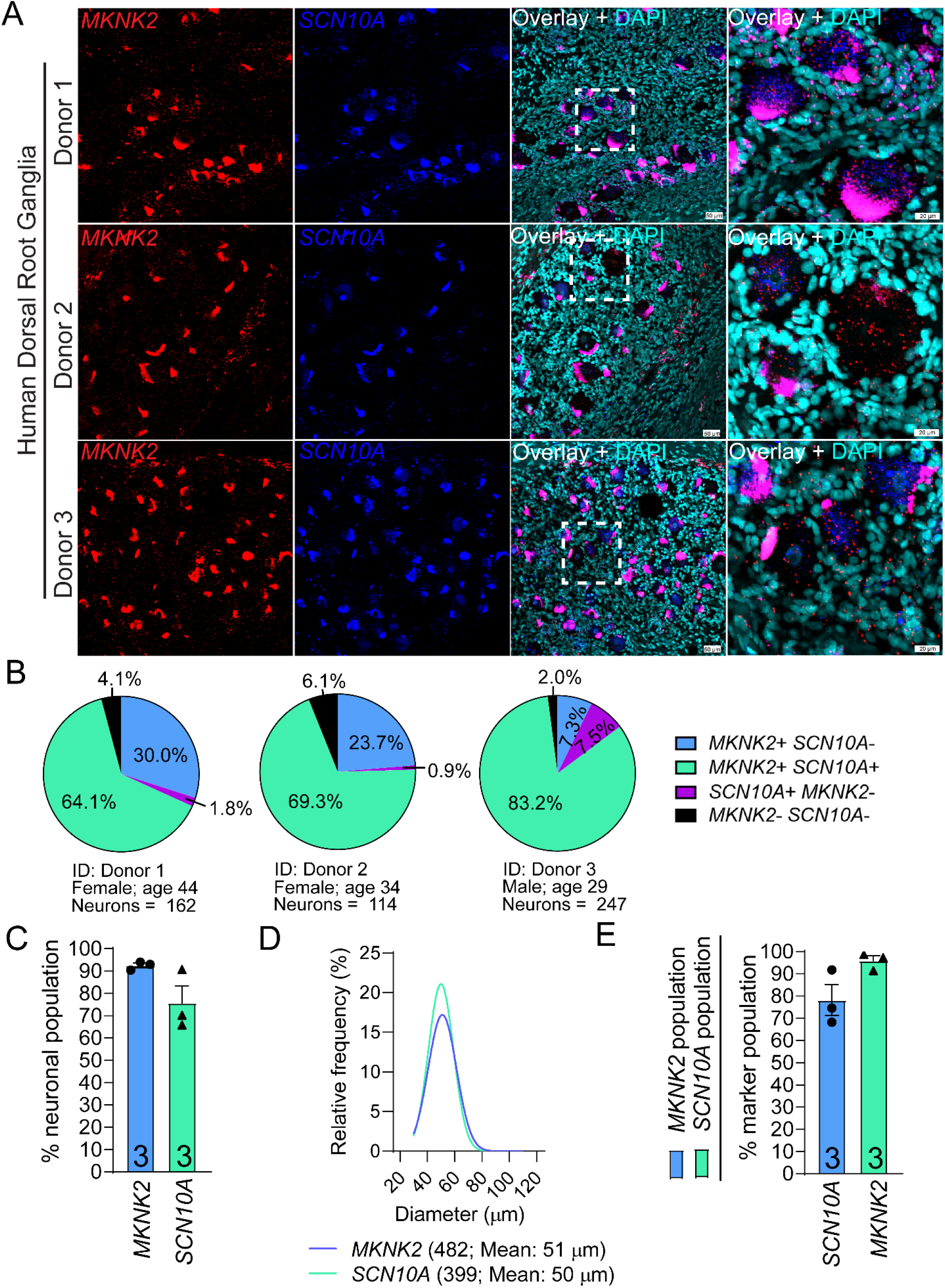
Distribution of *MKNK2* (MNK2) mRNA in human dorsal root ganglia. **A)** Representative 20X images of human lumbar DRGs labeled with RNAscope *in situ* hybridization for *MKNK2* (red) and *SCN10A* (blue) mRNAs and co-stained with DAPI (cyan). The fourth panel for each donor is a zoomed-in region demarcated by white boundaries in the 20X overlay image. Lipofuscin (globular structures) that autofluoresced in both channels and appear magenta in the overlay image were not analyzed as this is background signal that is present in all human nervous tissue. *MKNK2* mRNA was expressed in neurons and non-neuronal cells. **B)** Pie-charts showing the distribution of *MKNK2* neuronal subpopulations in human DRG for each donor. **C)** 92.5% of human DRG sensory neurons were positive for *MKNK2*. **D)** Histogram with Gaussian distribution displaying the size profile of all *MKNK2*+ and *SCN10A*+ neurons in human DRG. **E)** Percentage of *MKNK2*+ neurons that coexpressed *SCN10A* (blue bar), and the percentage of *SCN10A*+ neurons that coexpressed *MKNK2* (green bar). Scale bars: 20X = 50μm. Zoomed in panel = 20μm.

Next, we conducted RNAscope for *MKNK1* mRNA in combination with *SCN10A* on human TG to assess its cellular expression in human TG sensory neurons. We found that *MKNK1* mRNA was abundant in all human TG neurons (100%) (**Fig 3A-C**), and in non-neuronal cells as well. These neurons ranged in size from small-to-medium diameter (**Fig 3D**). 100% of *SCN10A*+ nociceptive neurons coexpressed *MKNK1* mRNA (**Fig 3E**).

**Figure 3.**
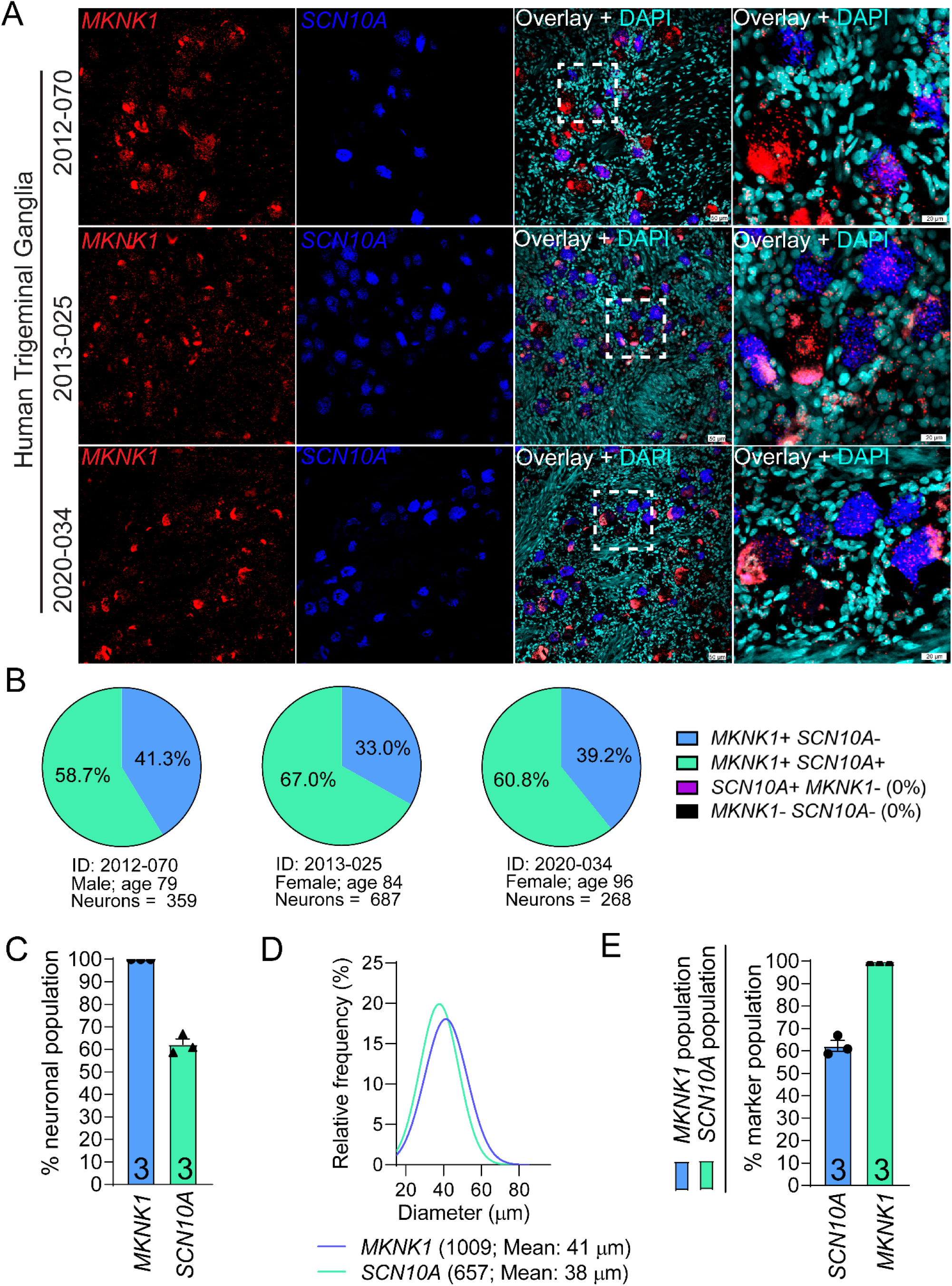
Distribution of *MKNK1* (MNK1) mRNA in human trigeminal ganglia. **A)** Representative 20X images of human TGs labeled with RNAscope *in situ* hybridization for *MKNK1* (red) and *SCN10A* (blue) mRNAs and co-stained with DAPI (cyan). The fourth panel for each donor is a zoomed-in region demarcated by white boundaries in the 20X overlay image. Lipofuscin (globular structures) were not analyzed as this is background signal that is present in all human nervous tissue. *MKNK1* mRNA was expressed in neurons and non-neuronal cells. **B)** Pie-charts showing the distribution of *MKNK1* neuronal subpopulations in human TG for each donor. **C)** 100% of human TG sensory neurons were positive for *MKNK1*. **D)** Histogram with Gaussian distribution displaying the size profile of all *MKNK1*+ and *SCN10A*+ neurons in human TG. **E)** Percentage of *MKNK1*+ neurons that coexpressed *SCN10A* (blue bar), and the percentage of *SCN10A*+ neurons that coexpressed *MKNK1* (green bar). Scale bars: 20X = 50μm. Zoomed in panel = 20μm.

Finally, we assessed the distribution of *MKNK2* mRNA in human TG. *MKNK2* mRNA was abundantly expressed in all neurons (100%), and qualitatively appeared to be more abundant than *MKNK1* in human TG (**Fig 4A-C**). It was present in non-neuronal cells as well. *MKNK2*-positive neurons ranged in size from small-to-medium diameter (**Fig 4D**). Like *MKNK1*, all SCN10A+ (100%) were copositive for MKNK2 (**Fig 4E**).

**Figure 4.**
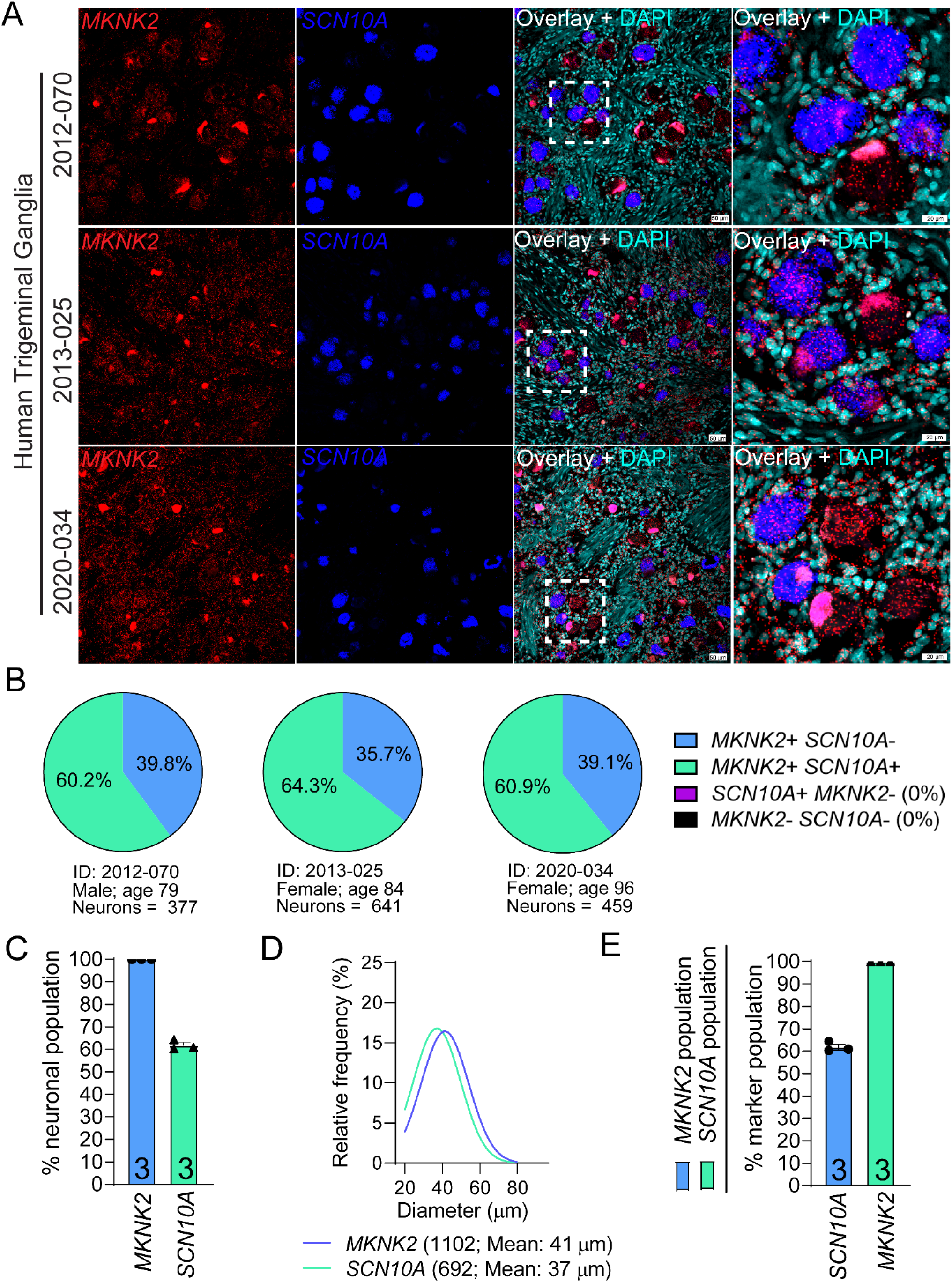
Distribution of *MKNK2* (MNK2) mRNA in human trigeminal ganglia. **A)** Representative 20X images of human TGs labeled with RNAscope *in situ* hybridization for *MKNK2* (red) and *SCN10A* (blue) mRNAs and co-stained with DAPI (cyan). The fourth panel for each donor is a zoomed-in region demarcated by white boundaries in the 20X overlay image. Lipofuscin (globular structures) were not analyzed as this is background signal that is present in all human nervous tissue. *MKNK2* mRNA was expressed in neurons and non-neuronal cells. **B)** Pie-charts showing the distribution of *MKNK2* neuronal subpopulations in human TG for each donor. **C)** 100% of human TG sensory neurons were positive for *MKNK2*. **D)** Histogram with Gaussian distribution displaying the size profile of all *MKNK2*+ and *SCN10A*+ neurons in human TG. **E)** Percentage of *MKNK2*+ neurons that coexpressed *SCN10A* (blue bar), and the percentage of *SCN10A*+ neurons that coexpressed *MKNK2* (green bar). Scale bars: 20X = 50μm. Zoomed in panel = 20μm.

We also observed *MKNK1* and *MKNK2* mRNA puncta in the fiber-rich areas of the human TG (**Fig 5**). While most of the signal appeared to be localized around non-neuronal nuclei, likely Schwann cells given their elongated shape and proximity to axons, there was some puncta that appeared to be localized within the axon. This signal is likely axonal-trafficked *MKNK1* (**Fig 5A-B**) and *MKNK2* mRNAs (**Fig 5C-D**). Qualitatively, there appeared to be more *MKNK2* than *MKNK1* mRNA localized within the axons (**Fig 5D**).

**Figure 5.**
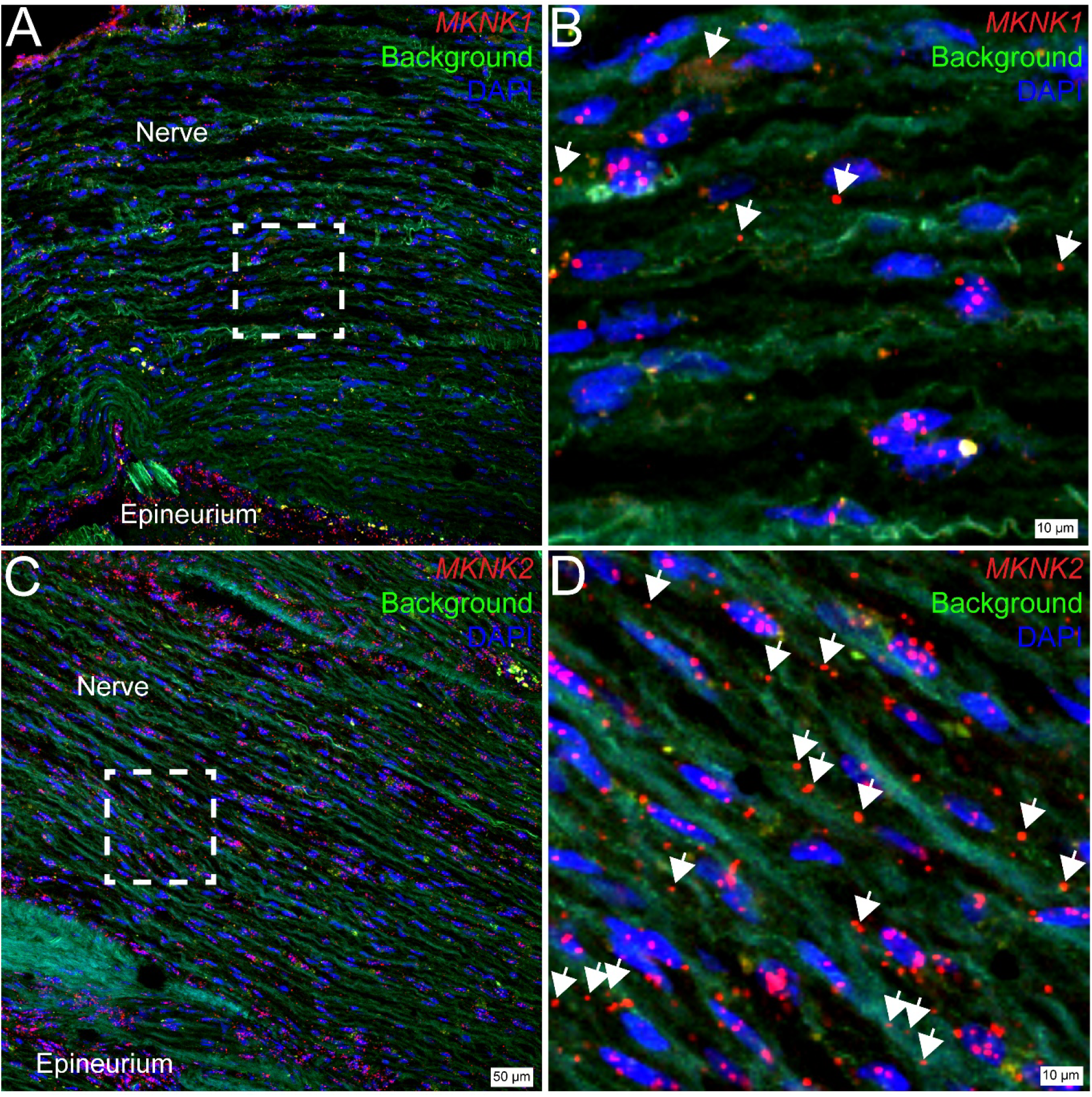
Axonal *MKNK1* (MNK1) and *MKNK2* (MNK2) mRNAs in the human trigeminal ganglia. **A)** Representative 20X image of the axonal-rich area of the human TG labeled with RNAscope *in situ* hybridization for *MKNK1* (red) mRNA and co-stained with DAPI (blue). Background autofluorescence was brightened to visualize the axons (green). **B)** Zoomed-in region of the nerve demarcated by white boundaries in panel A. *MKNK1* mRNA can be seen in or adjacent to non-neuronal nuclei. White
arrows point towards *MKNK1* mRNA puncta that are localized within the axon. **C)** Representative 20X image of the axonal-rich area of the human TG labeled with RNAscope *in situ* hybridization for *MKNK2* (red) mRNA and co-stained with DAPI (blue). Background autofluorescence was brightened to visualize the axons (green). **D)** Zoomed-in region of the nerve demarcated by white boundaries in panel C. *MKNK2* mRNA can be seen in or adjacent to non-neuronal nuclei. White arrows point towards *MKNK2* mRNA puncta that are localized within the axon. Scale bars: 20X = 50μm. Zoomed in panel = 10μm.

## Discussion

The primary conclusion we reach based on these experiments is that both *MKNK1* and *MKNK2* genes are expressed in human nociceptors in the DRG and TG. Given the very high percentage of neurons that co-expressed *SCN10A* for both MNK genes, it is likely that nearly all human nociceptors express both isoforms at the mRNA level. Mouse single nucleus RNA sequencing experiments likely underestimate gene expression for both genes due to under-sampling by only assessing recently transcribed genes (Stark et al., 2019), however, our results still suggest a potential species difference in isoform expression. In mice, *Mknk1* is more highly expressed by peptidergic nociceptors and *Mknk2* more highly expressed by non-peptidergic nociceptors (Zeisel et al., 2018; Renthal et al., 2020) whereas in humans both isoforms are expressed across nociceptor subtypes with qualitatively higher expression of *MKNK2* in most neurons, especially those residing in the TG.

In mice, *Mknk1* and *Mknk2* double knockout leads to reduced inflammatory and neuropathic pain and a nearly completely loss of eIF4E phosphorylation in the DRG (Moy et al., 2017). *Mknk1* knockout recapitulates the phenotype of the double knockout even though only about 50% of eIF4E phosphorylation is lost in the DRG (Moy et al., 2017). This suggests that the *Mknk1* isoform may play a more important role in mouse pain models. While one MNK1-specific inhibitor has been described (Santag et al., 2017), it has not been tested in mouse pain models. A specific MNK1/2 inhibitor, eFT508 (Reich et al., 2018), has been tested in multiple mouse pain models and this compound produces an anti-nociceptive effect that is similar to what is seen in Mknk1 knockout mice (Megat et al., 2019; Shiers et al., 2020b). eFT508 nearly completely eliminates eIF4E phosphorylation in the mouse DRG and brain when it is given systemically at doses that are anti-nociceptive (Megat et al., 2019; Shiers et al., 2020b). Based on these previous mouse studies, one might conclude that MNK1 is the more important isoform to target in pain clinical trials. Our human results suggest caution in relying on this interpretation of the mouse data because species differences in gene expression in nociceptor subtypes may lead to a more prominent role for MNK2 in the sensitization of human nociceptors.

We used two different RNAscope kits in our experiments, the V1 kit for DRG and V2 kit for TG. DRG experiments were done before the V2 kit was widely adopted for use and the TG experiments were done in late 2022, after the V1 kit had been discontinued and the V2 kit was more widely used, and we had carefully characterized it for use on human neuronal tissues. We observed a better signal to noise ratio for the V2 kit, and this likely explains why a slightly larger proportion of neurons in the human TG were positive for both *MKNK* genes. We think that the TG results likely more accurately reflect expression for both *MKNK* isoforms and that lack of detection in some nociceptors in human DRG is due to less sensitivity of signal for the V1 kit. This raises a question of whether MNK1 and MNK2 are ubiquitously expressed in all cells in the body. eIF4E is a ubiquitously expressed gene, and can presumably be phosphorylated in all cells, a process that likely depends on expression of at least one MNK isoform. Single cell sequencing efforts across cell types in mouse (Tabula Muris et al., 2018) and human (Tabula Sapiens et al., 2022) show that *MKNK1* and *MKNK2* are uniquely expressed in certain cell types. Our results show clearly that human nociceptors express both *MKNK* genes in the same cell.

An unexpected finding in our experiments was the strong expression of *MKNK1* and *MKNK2* mRNA in axonal fields in the human TG. Some of this mRNA signal is likely due to expression in Schwann cells but given the low expression level for both *Mknk* genes in mouse peripheral nerve single nucleus datasets, this is unlikely to explain all of the signal (Brosius Lutz et al., 2022; Yim et al., 2022). Another explanation is transport of *MKNK1* and *MKNK2* mRNA into the axons of human sensory neurons. This is an interesting possibility given that MNK1/2 is proposed to control the local, activity-dependent translation of mRNA that are important for the sensitization of nociceptor axons in response to local cues (Yousuf et al., 2021). While further work will be needed to understand the dynamics of localization of *MKNK1* and *MKNK2* mRNAs in human sensory neuronal axons, this finding is consistent with the important role already described for these genes in regulating nociceptor excitability based on mouse studies.

We conclude that dual MNK1 and MNK2 inhibitors should be developed for testing as a novel treatment for pain based on expression of both genes in nearly all human nociceptors. Dual MNK inhibitors have progressed through phase II clinical trials for oncology and are well tolerated except for dose-limiting CNS-mediated side effects (Falchook et al., 2017; El-Khoueiry et al., 2020). This clinical data suggests that dual MNK inhibitors for pain can be progressed to clinical testing with appropriate optimization to increase the safety window for these inhibitors.

## Acknowledgments

This work was supported by NIH grants NS065926 and NS115692. The authors are grateful to the organ donors and their families for the gift of life and research provided by their organ donation. The authors thank Anna Cervantes and Geoffrey Funk of the Southwest Transplant Alliance for tissue recovery.

